# Short-term memory capacity and chronic stress levels predict cognitive effort choice as a function of reward level and effort demand

**DOI:** 10.1101/2025.07.24.666659

**Authors:** Brandon J. Forys, Catharine A. Winstanley, Rebecca M. Todd

## Abstract

Every day, we make choices about how much effort we are willing and able to use to achieve the outcomes we desire against the backdrop of constantly shifting effort demands and available rewards. While factors like visual short-term memory and chronic stress levels can predict responses to stable cognitive effort demands, we do not yet know whether they constrain one’s choices of higher effort trials for larger rewards when task demands and potential outcomes shift over time. Here, we examined whether these factors predicted the choice to deploy cognitive effort given increasing effort demands and the tendency to deploy effort given shifting reward availability. Undergraduate participants first performed an online visual short-term memory task to assess capacity for visuospatial short-term memory. They then completed a series of choice trials where they could choose between high-effort, high-reward or low-effort, low-reward trials. In two blocks, we varied either the effort required on high-effort trials or the reward offered on both trial types. We found that visual short-term memory predicted the likelihood of choosing high-effort trials given shifting rewards, while chronic stress and everyday preferences for cognitively effortful strategies predicted the tendency to deploy increasing amounts of effort for a stable reward. Furthermore, participants’ subjective reports show a strong focus on attentional processes, and balancing rewards and losses, when making decisions about how much effort to deploy. These findings shed light on distinct trait-level factors associated with cognitive effort choices given shifting demands and outcomes.

**Significance statement:** We must often choose how much work to put in to complete everyday tasks. However, we do not know what behavioural factors drive these choices in humans when the effort required to complete a task - or potential rewards - shifts over time. In a visual short-term memory task adapted from rodent work, we found that those with higher visual short-term memory ability chose more high effort trials as effort demands increased, while chronic stress and everyday preferences for effortful strategies predicted more effort for a reward. Furthermore, participants described prioritizing sustaining attention in order to successfully complete the task. These findings shed light on distinct trait-level factors associated with cognitive effort choices given shifting demands and outcomes.

## Introduction

We must often deploy sustained attention to meet everyday task demands. For example, when writing a report, we must choose how much time to spend on it to meet our goals without overworking ourselves. These choices require cognitive control deployment through cognitive effort. There is growing interest in studying what drives effortful choices (Shenhav et al., 2013, 2017; Bustamante et al., 2023), comparing the value of potential rewards to the cost of effort deployment (Apps et al., 2015; Westbrook and Braver, 2015) and discounting further effort if needed (Westbrook et al., 2013). However, crucial questions remain about the behavioural and environmental factors that influence one’s choices in response to these effort demands.

Past studies have identified individual differences in cognitive capacity that can influence effort deployment (Grahek et al., 2019). In humans, visual short-term memory ability predicted choices of higher rewards given static effort demands in a memory task (Forys et al., 2024), echoing findings from rodent tasks showing higher preferences for high-effort trials among better-performing rodents (Hosking et al., 2016; Hales et al., 2024). Higher chronic stress and depression levels are associated with suboptimal strategies (Lenow et al., 2017; Horne et al., 2021) and impairments in executive functions such as working memory (Watt et al., 2017). However, real-world cognitive effort demands and corresponding rewards often shift over time (Westbrook et al., 2013; Gratton et al., 2018). For example, a video game may differ in difficulty and rewards as the player gains experience over time. Delay discounting - the desire to trade off additional rewards given longer waits - also factors into valuations of effort demands (Massar et al., 2020; Toro-Serey et al., 2022). Thus, we do not yet understand the impacts of these individual differences in chronic stress and depression levels on our engagement with changing task difficulties.

Choices when evaluating shifting effort demands can also be influenced by contextual or trait level constraining factors related to effort benefit and demand signals (Bustamante et al., 2024). These include trait-level factors of individuals reflecting everyday effort attitudes (Sandra and Otto, 2018). The Need for Cognition scale (NCS; Wu et al. (2014)) probes individual differences in willingness to seek out challenging tasks. Westbrook et al. (2013) found that low NCS scores were associated with fewer choices of high effort, high-reward trials and higher points of subjective equivalence where low and high effort trials were equally preferred. However, studies examining in-lab effort choices typically do not address questions of whether task performance reflects real-life choice behaviour.

To probe whether choices to deploy cognitive effort elicited by laboratory tasks reflect choices made outside the lab, one must query participants’ experiences to examine whether they capture everyday approaches to cognitive effort. Subjective assessments of choice behaviours (Milyavskaya et al., 2021; Bustamante et al., 2023), through methods like subjective report (Braun and Clarke, 2006), can indicate how people make real-world effort deployment decisions. However, no study has yet looked at whether choices about shifting effort demands capture everyday cognitive effort.

To answer these outstanding questions, in the present study we 1) examined the degree to which individual differences in visual short-term memory ability, chronic stress, and NCS scores were associated with effort choices when effort demands or rewards shift, and 2) documented participant reports of the applicability of in-lab effort demands to everyday tasks. To address these goals, we presented a novel cognitive effort task that simultaneously manipulated both effort demands and rewards outcomes - built upon the work of Forys et al. (2024) and Luck and Vogel (1997), and adapted from the rodent Cognitive Effort Task (rCET; Hosking et al. (2016)) for translational validity. Participants chose between high effort, high reward (HR) trials or low effort, low reward (LR) trials, experiencing shifting effort demands or reward levels, before providing written responses to questions about task experiences. We evaluated individual differences in short-term memory ability, chronic stress levels, depression levels, and NCS scores. Furthermore, through a thematic analysis approach, we examined themes that characterized the strategies and information that participants drew on during the task and in comparable real-life scenarios from subjective reports. From Forys et al. (2024)’s findings that visual short-term memory predicted overall increases of high effort trials, we predicted that higher visual short-term memory ability would drive increased preferences for high effort trials despite increasing effort demands. Furthermore, we predicted that higher chronic stress and depression levels would be associated with decreased high effort trial preferences given diminishing rewards (Bishop and Gagne, 2018). From Westbrook et al. (2013) and Crawford et al. (2021), we predicted that lower NCS scores would drive increased discounting of high effort options.

## Materials and Methods

As in Forys et al. (2024), we powered our study through the pwrss package (Bulus, 2023) to achieve an expected power of 80% at an odds ratio of 0.70 for a logistic regression evaluating the probability of a participant selecting 70% or more high effort, high reward trials (being in the high effort group in the study), giving us a target sample size of *N* = 487. We recruited *N* = 527 participants from the Human Subject Pool of psychology undergraduate students at the University of British Columbia; they were each compensated with bonus points for their courses as well as a CAD $5.00 Starbucks gift card. Of these, *n* = 80 participants did not complete the initial survey. Furthermore, *n* = 6 participants spent more than 30 seconds choosing any one trial - these participants were excluded as this level of choice latency could indicate disengagement with task demands. We also excluded *n* = 6 participants with K estimate scores on set size = 4 that were over 3 standard deviations from the mean. Here, the whole-display K estimate score (Pashler, 1988) for set size = 4 captures participant’s visual short term memory ability as the hit rate minus the false alarm rate - divided by 1 minus false alarm rate - when participants had to encode 4 shapes, which was determined in Forys et al. (2024) to be the largest set size on which all participants could reliably complete the task. In total, we analyzed data from *N* = 413 participants (*n* = 61 male, *n* = 339 female, *n* = 13 other; Table 1). The study was approved by the Behavioural Research Ethics Board at the University of British Columbia, approval code H20-01388.

**Table 1:** Demographic information for all participants, by sex and effort deployment group. BDI prop. = Beck Depression Inventory II proportion score (score divided by max score). PSS = Perceived Stress Scale score. DARS = Dimensional Anhedonia Rating Scale. NCS = Need for Cognition Scale score. High effort group: > 70% HR trials selected; low effort group: <= 70% HR trials selected.

| Group | Sex | n | M<sub>age</sub> | SD<sub>age</sub> | Min<sub>age</sub> | Max<sub>age</sub> | M<sub>BDIprop</sub> | SD<sub>BDIprop</sub> | M<sub>PSS</sub> | SD<sub>PSS</sub> | M<sub>NCS</sub> | SD<sub>NCS</sub> |
| --- | --- | --- | --- | --- | --- | --- | --- | --- | --- | --- | --- | --- |
| High effort | Female | 256 | 20.36 | 2.62 | 12 | 46 | 0.25 | 0.18 | 19.86 | 6.14 | 57.40 | 12.46 |
|  | Male | 50 | 21.72 | 5.36 | 18 | 49 | 0.23 | 0.17 | 17.96 | 5.52 | 59.89 | 17.26 |
|  | Other | 10 | 21.00 | 2.35 | 17 | 26 | 0.20 | 0.17 | 18.50 | 2.64 | 58.67 | 8.00 |
| Low effort | Female | 83 | 20.06 | 1.59 | 17 | 26 | 0.31 | 0.18 | 21.92 | 6.26 | 52.91 | 14.19 |
|  | Male | 11 | 21.00 | 2.14 | 19 | 25 | 0.20 | 0.17 | 15.09 | 6.69 | 57.17 | 12.89 |
|  | Other | 3 | 18.67 | 0.58 | 18 | 19 | 0.15 | 0.18 | 17.00 | 7.00 | 60.00 | NA |

### Stimulus Presentation

All participants completed the study online on their own devices, via the Pavlovia online study platform using PsychoPy 2024.1.5 (RRID: SCR_006571) (Peirce et al., 2019). As the study required keyboard responses, participants were not allowed to complete the study on mobile phones or tablets - devices on which the onscreen keyboard could block one’s view of the stimuli.

All stimuli used in the study were implemented in PsychoPy (Peirce et al., 2019) and were adapted from Forys et al. (2024). As a measure of each individual’s visuospatial short-term memory capacity, we presented a visuospatial short-term memory task (Fig. 1A, B) modified from Luck and Vogel (1997) and adapted from an open-source version of the task on Pavlovia (de Oliveira, João Roberto Ventura, 2023). We presented a series of trials with between 2 and 8 coloured squares that were presented for 500 ms. Each square subtended a visual angle of approximately 0.05*^◦^* on the screen. A multi-coloured square mask would then appear at each of the original squares’ locations for another 500 ms, followed by a single coloured square appearing in one of the positions of the original squares. This final square had a 50% chance of being the same or a different colour from the square appearing in the same position in the initial part of the trial. After indicating with a keyboard press whether the square was the same or a different colour from the initial square, they would see whether or not they gained points towards a monetary reward.

**Figure 1:**
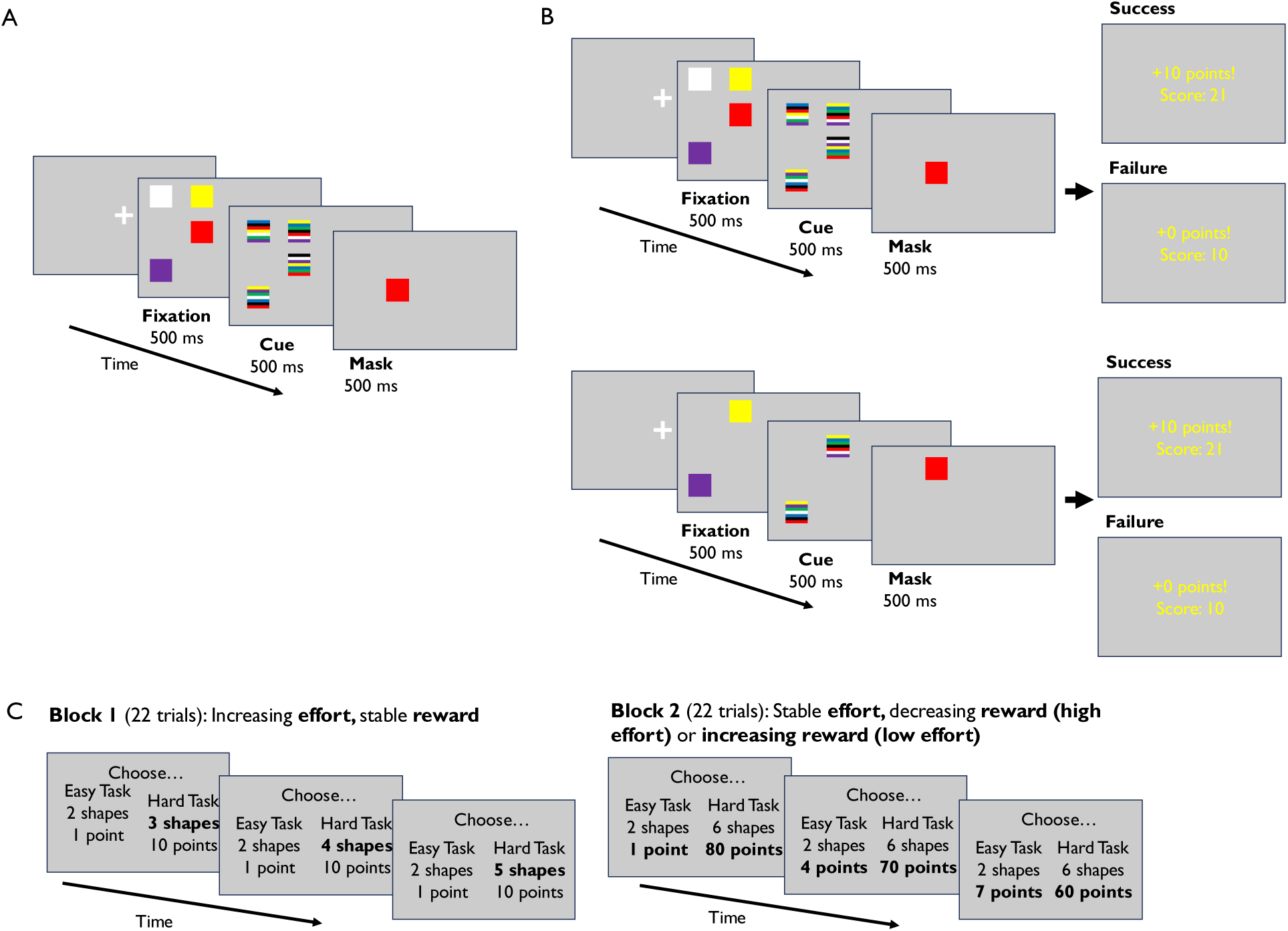
Trial layout diagram. Layout of the experimental tasks. (A) The calibration phase of the change detection task, where participants saw an array of either 2, 4, 6, or 8 squares and indicated whether the final square that appeared (the probe) was of the same or a different colour from the square that appeared in the same location in the previously show array. (B, C) The reward phase of the change detection task, in which participants selected either low effort, low reward or high effort, high reward trials in two blocks. In Block 1, effort demands increased while rewards remained fixed. In Block 2, rewards decreased on high effort trials and increased on low effort trials while effort demands remained fixed. After making their choice, once again, participants indicated with a keyboard press whether the final square that appeared (the probe) is the same or a different colour from the square that appeared in the same location in the array. Here, if they made the correct decision, in Block 1 they would receive 1 point (low effort trial) or 10 points (high effort trial) while in Block 2 they would receive a variable amount of reward. In all cases, they would receive 0 points for an incorrect decision. The order of blocks was counterbalanced and the progression of increasing effort demands or shifting rewards was randomized within each block. Factors that change between trials are bolded for emphasis here, but were not bolded in the actual stimuli.

### Procedure

#### Questionnaires

Participants began the study by completing an online questionnaire. After giving consent and demographic information, they were asked whether they had a history of depression and anxiety and, if so, whether they received treatment or medication for these conditions. They were then given the Perceived Stress Scale (PSS; Levenstein et al. (1993)) - an index of chronic stress; the Beck Depression Inventory II (BDI; Beck et al. (1996)) - a clinically validated and reliable (Richter et al., 1998) index of depression; the Dimensional Anhedonia Rating Scale (DARS; Rizvi et al. (2015)) - a reliable index of anhedonia, or impairments to one’s pleasure response, that has been validated in a variety of populations (Arrua-Duarte et al., 2019; Gorostowicz et al., 2023); and the Need for Cognition Scale (NCS; Cacioppo and Petty (1982)) - an index of everyday attitudes towards completing cognitively effortful tasks showing high test-retest reliability (Sadowski and Gulgoz, 1992). Afterwards, they were redirected to the first phase of our study, the visuospatial short-term memory task.

#### Experimental design

As in Forys et al. (2024), we used an adaptation of a visual short term memory k-estimate task (Luck and Vogel, 1997) for two purposes. In the first, ***assessment phase*** of the experiment, this task served to measure individual differences in visuospatial short-term memory, providing a single score for visual short term memory capacity to be used as a predictor in analyses of choice behaviour on the subsequent choice task. Visual short term memory is important for incorporating and manipulating information about task demands to complete everyday tasks; therefore, measuring this construct provides a useful index of cognitive effort ability. After receiving instructions about which stimuli would appear and how to respond to them, participants first completed a series of ten practice trials presented in a random order. On half of these trials, the probe square was the same colour as that of the cue square (congruent/no change trial); on the other half, the probe square was a different colour from that of the cue square (incongruent/change trial). Of these practice trials, four trials had a set size = 2, another four trials had a set size = 4, and another two trials had a set size = 6. In each trial, the probe square could be anywhere in the array, and participants were required to hold the whole array in mind in order to successfully indicate whether the probe color had changed. Following these practice trials, participants completed 40 trials of the k-estimate task with 20 change and 20 no change trials (Fig. 1B). In total, 10 trials of each set size (2, 4, 6, and 8 squares onscreen) were presented in a randomized order. Although the original task described in Luck and Vogel (1997) contained trials with up to 10 squares onscreen, this largest set size was determined to be too difficult for participants to reliably complete correctly following piloting; as such, the maximum set size was 8.

Furthermore, although the main task had a maximum set size of 8, the practice block only included a maximum set size of 6 so as to give a brief overview of the trials and required responses without giving excessive practice with a large set size, for which performance would be evaluated in the subsequent block. Participants did not receive feedback on whether their responses were correct or not in the practice block or the memory task, in keeping with Luck and Vogel (1997).

The primary dependent variable of interest in this study was choice of high effort vs. low effort trials while we independently manipulated the amount of cognitive effort (operationalized as set size) and reward. In the second, ***choice phase*** of the experiment we used a task (Fig. 1C) adapted from that used in Forys et al. (2024) which, in turn, was adapted from a rodent cognitive effort task (rCET) (Hosking et al., 2016; Silveira et al., 2021). In each trial, participants had to choose between two k-estimate sets requiring varying levels of cognitive effort/offering varying levels of reward. Choices were presented in two counterbalanced blocks manipulating effort demands and potential rewards. In the *effort increase* block, the set size offered in the high effort trials increased by 1 at a normally distributed interval of every 3 trials on average, such that the high effort set size offered would increase from 3 to 10. Crucially, these increases were randomly ordered across the block, such that the overall effort increase was non-sequential - allowing changes in performance and effort preferences to be disentangled from potential fatigue induced by linearly increasing effort demands. The low effort trials always displayed a set size of 2, and the reward levels remained constant across the block for both low and high effort trials at 1 point for low effort trials and 10 points for high effort trials. Thus, the amount of cognitive effort required to successfully achieve the higher reward varied over time in the effort increase block.

In the *shifting reward* block, the reward offered on high effort trials decreased by 10 from a maximum of 80 points to a minimum of 10 points at a normally distributed interval of every 3 trials on average. Conversely, the reward offered on low effort trials increased by 3 points from a minimum of 1 point to a maximum of 22 points at a normally distributed interval of every 3 trials on average. Thus, the reward levels offered converged between low and high effort trials. High effort and low effort demands remained constant throughout this block at set sizes of 6 and 2, respectively. This level of difficulty was previously determined to offer a moderate level of challenge to all participants (Forys et al., 2024). As in the increasing effort block, these reward changes were randomly ordered; however, combinations of low and high effort reward amounts always remained the same (i.e. a trial offering 70 points on a high effort trial would always be paired with a trial offering 4 points on a low effort trial). Because of the normal distribution of effort and reward intervals - to prevent a strategy of purposely shifting choices at every fixed interval of trials - not every participant saw all point or reward intervals in a given block.

Following the choice phase of the experiment, participants answered, in written form, a series of three questions about their effort choice preferences, cognitive effort strategies, and the extent to which the task’s effort demands were relevant to real-life effort demands. Responses to these questions allowed us to document the ecological validity of our measure in terms of how it reflected real-world decisions about shifting effort demands and outcomes.

#### Statistical analysis

All analyses were conducted using R 4.4.3 “Trophy Case” (R Development Core Team, 2011) through RStudio (Booth et al., 2018).

Our primary question of interest was whether preferences for high effort trials were significantly predicted by visual short term memory ability, chronic stress levels, or Need for Cognition scores given increasing effort demands or shifting reward levels. As in Forys et al. (2024), the primary predictor variables in our study were 1) visual short term memory ability, operationalized as a participant’s K estimate at a set size of 4 in the first, *assessment phase* of the experiment; 2) depressive traits, operationalized as a participant’s BDI proportion score; and 3) chronic stress levels, operationalized as a participant’s PSS score. Additionally, to evaluate overall preferences for deploying cognitive effort strategies, we measured 4) everyday cognitive effort preferences, operationalized as a participant’s score on the Need for Cognition scale (NCS; Cacioppo and Petty (1982)). The primary dependent variable in our study was 1) high effort preference, operationalized as the proportion of High Effort, High Reward (HR) trials selected in each block of the task (increasing effort or shifting reward). Furthermore, in support of this analysis, we investigated 2) accuracy, operationalized as the proportion of correct responses (hits and correct rejections), and 3) choice latency, operationalized as the time in seconds until participants chose the high or low effort option for each trial. We compared high effort preference, accuracy, and choice latency between increasing effort and shifting reward blocks.

We first divided participants into high and low effort preference groups for translational comparability with existing rodent and human work. As in Cocker et al. (2012) and Forys et al. (2024), participants who chose the HR option for more than 70% of all trials in the reward phase of the task comprised the high effort preference group, while those who chose the HR option for 70% or less of all trials in this phase comprised the low effort preference group. In the *choice phase* of the experiment, we varied either the effort level (set size) of visual short-term memory task trials or reward amounts in two blocks. We conducted three binomial logistic regressions to determine whether visual short term memory ability, chronic stress levels, or Need for Cognition score significantly predicted whether a participant was in the high or low effort preference group in terms of their effort choices in each block of the *choice task*. As choices for high-effort trials may could also vary with visual short term memory ability, we then conducted a linear regression to evaluate whether the above regressors also predict the proportion of high effort trials selected.

In addition to effort choice, we examined accuracy on each trial and choice latency for selecting the trial difficulty to evaluate whether participants were matched for performance and time spent selecting a trial (Cocker et al., 2012), regardless of how many high vs. low effort trials they selected. We ran two multi-level models - one for each block - through the lmerTest package (Kuznetsova et al., 2017) to evaluate whether sex, short term memory ability, depressive traits, chronic stress levels, Need for Cognition score, and effort preference group predicted choice latency or accuracy. Additionally, to assess the point of equivalence at which participants equally selected high vs. low effort trials, we examined the proportion of high effort trials chosen when each effort level or reward amount was offered. Here, we used a one-sample t-test to determine whether this proportion significantly differed from chance (50%).

Last, we examined patterns of responses to questions about the effort strategies used in the task - and their relevance to everyday task demands. Thematic analysis approaches (Braun and Clarke, 2013) are an effective method for capturing categories of information that participants have identified as being important to their experiences. However, conducting a manual thematic analysis on very large datasets can be highly time consuming. Large language models like ChatGPT or Microsoft Copilot have been shown to be effective at rapidly describing and extracting themes from large sets of text (Martínez-Pernía et al., 2025), like those produced in our study, with near-human levels of accuracy. Thus, we used Microsoft Copilot (based on GPT-4o) to evaluate each of the three questions we asked our participants, and identify categories and subcategories present in their responses, using the following prompt (adapted from Martínez-Pernía et al. (2025)):

*“The attached file contains participants’ responses to the question: [Question text] Create a 6-column table to provide a detailed description (bodily sensations, emotional states, intentions, attention, and cognitions) of the participants’ responses. Analyze each row of the participants’ responses. In the table, note down the participant number in the first column; write a category of experience in the second column and a subcategory of experience in the third column. In the fourth column, add a short phrase that describes the content. In the fifth column, include an example quote from the participant’s responses. In the sixth column, add a definition of the subcategory. Export this table to a .csv file and a Word document. Use the following column names: Participant, Category, Subcategory, Description Phrase, Example Quote, Definition. Make the category and subcategory names as consistent as possible with your previous responses about the previous file I uploaded.”*

The output data produced a set of categories and subcategories characterizing each participant’s response to each question. Upon manually examining the output data for adherence to thematic analysis standards and consistency of categorization across the three questions, we illustrate the frequency of each thematic category and subcategory for each question, and provide summary table with definitions for each category and subcategory using the following prompt:

*“Based on the attached table and the previously given prompts, create a table with 3 columns (Category, Subcategory, and Conceptual definition) summarizing each category and subcategory in the attached table and creating a conceptual definition based on the category names and the example quotes given. Export this table to a CSV file and a Word document.”*

## Results

### Depressive traits, chronic stress levels, and Need for Cognition score

We report summary statistics for depressive traits (BDI), chronic stress levels (PSS), and Need for Cognition Scale (NCS) scores in Table 1. After correcting for multiple comparisons using the Bonferroni method, women had significantly higher depressive trait scores (*t*(1,365.40) = 7.89, *p* = <0.001, *d* = 0.27) (Fig. 2A) and chronic stress scores (*t*(1,384.55) = 15.04, *p* <0.001, *d* = 0.50) (Fig. 2B), as well as lower Need for Cognition (NCS) scores (*t*(959.71) = -5.64, *p* <0.001, *d* = 0.50), than men (Fig. 2C). Levels of depressive traits (BDI) and chronic stress (PSS) were significantly correlated with each other; however, none of these measures were significantly correlated with visual short term memory ability (K estimate) or Need for Cognition score (NCS) (Fig. 3).

**Figure 2:**
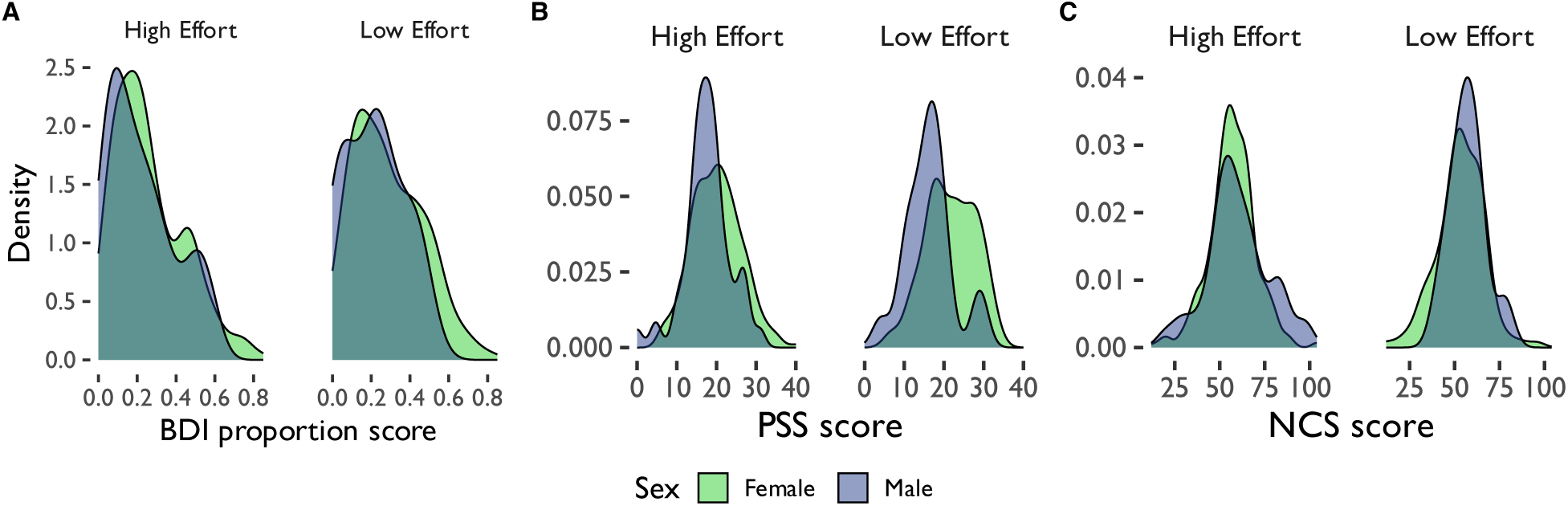
Demographic distributions. Distribution of (A) depressive trait (BDI) proportion scores (score divided by total possible score), (B) perceived stress (PSS) scores, and (C) Need for Cognition Scale (NCS) scores by sex and effort deployment group. BDI = Beck Depression Inventory, PSS = Perceived Stress Scale, NCS = Need for Cognition Scale.

**Figure 3:**
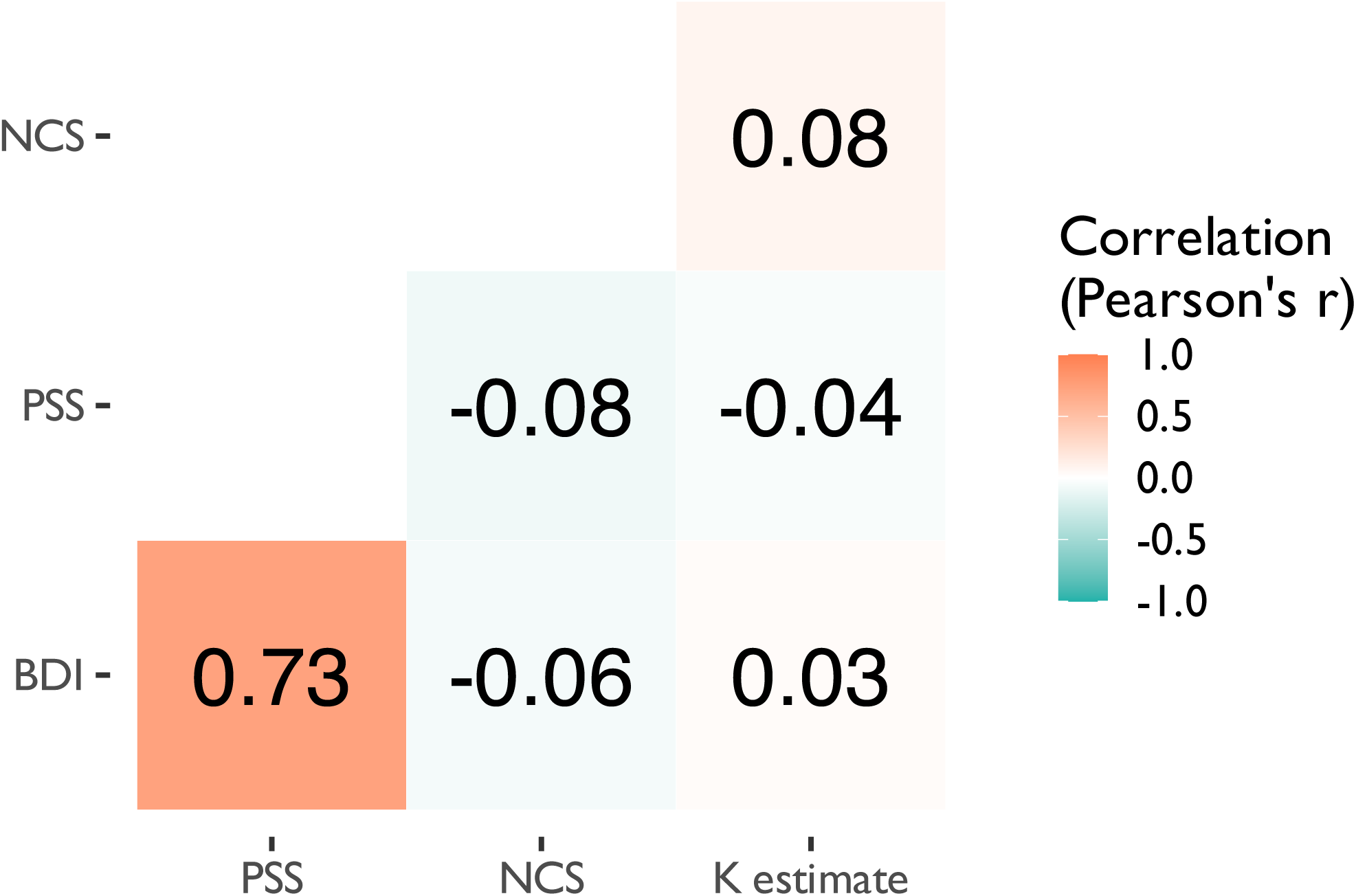
Binomial logistic regression predicting whether participants were in the high effort vs. low effort group from (A, B) visual short term memory ability; (C, D) chronic stress (PSS) level; and (E, F) Need for Cognition (NCS) score in the (A, C, E) increasing effort or (B, D, F) shifting reward blocks. Binomial logistic regression predicting whether participants were in the high effort vs. low effort group from (A, B) visual short term memory ability; (C, D) chronic stress (PSS) level; and (E, F) Need for Cognition (NCS) score in the (A, C, E) increasing effort or (B, D, F) shifting reward blocks. High effort group: > 70\% HR trials selected; low effort group: <= 70\% HR trials selected.

### Predictors of overall high vs. low effort choices

We were primarily interested in understanding what factors predict the choice of high vs. low effort/reward options given increasing effort demands vs. shifting reward levels (Table 2; Fig. 4). Our binomial regression showed that visual short term memory (K estimate) scores significantly predicted whether a participant was in the high effort or the low effort group in the *reward shift* block only (*z* = 2.52, *p* = 0.012, *e^β^*= 1.24). In contrast, chronic stress (PSS) significantly predicted whether a participant was in the high effort or the low effort group in the *effort increase* block only (*z* = -2.05, *p* = 0.04, *e^β^* = 0.96), as did Need for Cognition (NCS) score (*z* = 2.51, *p* = 0.012, *e^β^* = 1.03). Participants with higher visual short term memory ability had a higher probability of being in the high effort group, but only when reward levels were manipulated. On the other hand, participants with lower chronic stress levels and higher Need for Cognition scores had a higher probability of being in the high effort group, but only when effort demands increased. Depressive traits did not significantly predict whether participants were in the low vs. high effort groups. Furthermore, a linear model analysis with the same predictors as well as sex revealed that, across both blocks, only chronic stress, Need for Cognition scores, and sex significantly predicted the proportion of high effort trials that a participant selected (probability scores provided in Table 3).

**Figure 4:**
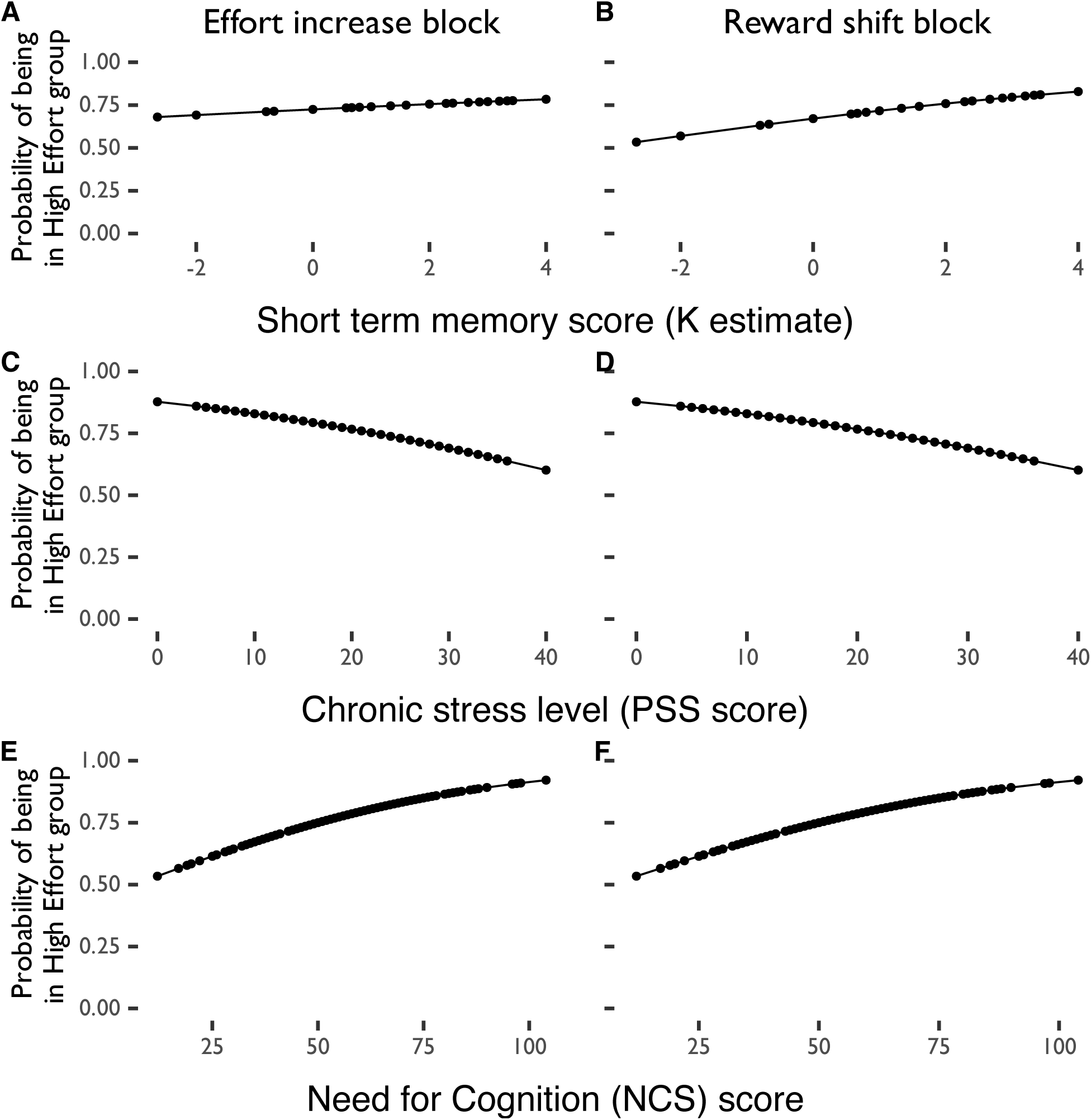
Binomial logistic regression predicting whether participants were in the high effort vs. low effortgroup from (A, B) visual short term memory ability; (C, D) chronic stress (PSS) level; and (E, F)Need for Cognition (NCS) score in the (A, C, E) increasing effort or (B, D, F) shifting rewardblocks. High effort group: > 70% HR trials selected; low effort group: <= 70% HR trials selected.

**Table 2:**
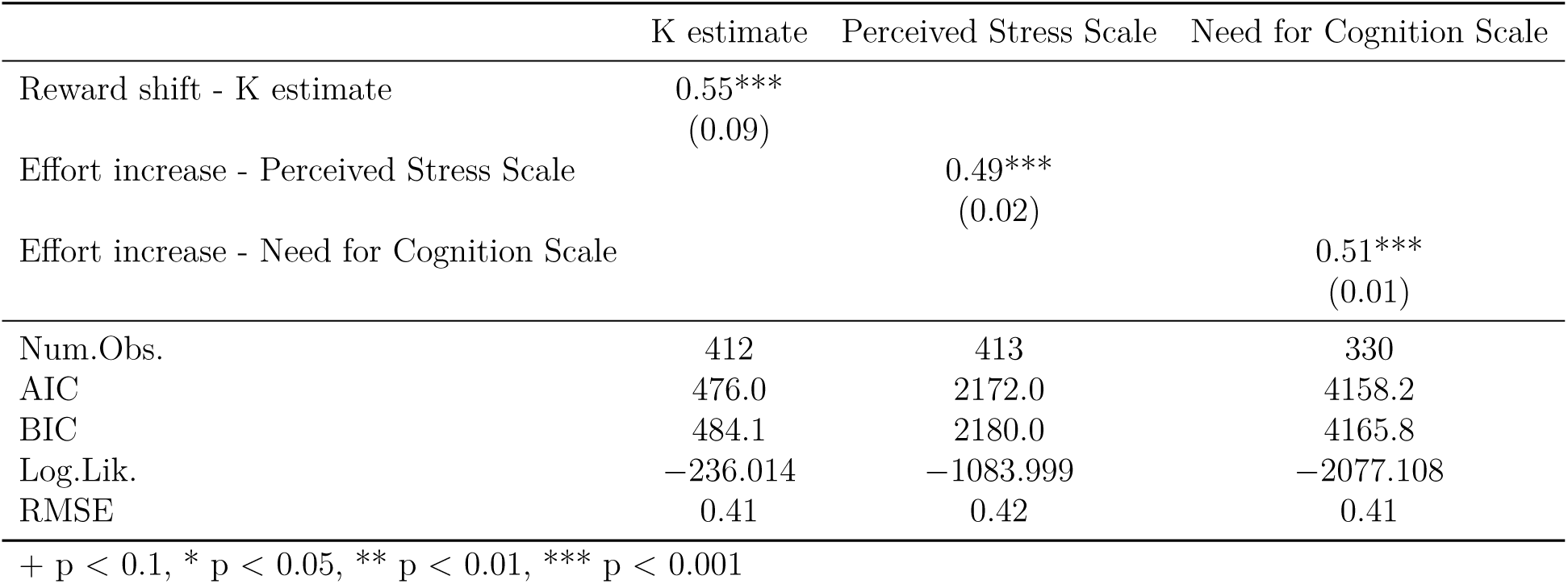
Binomial logistic regressions predicting whether participants were in the high effort vs. low effort group across both task blocks. All logits have been transformed to probability values. High effort group: > 70% HR trials selected; low effort group: <= 70% HR trials selected. K estimate = visual short term memory ability. AIC = Akaike information criterion, BIC = Bayesian information criterion, ICC = intra class correlation, RMSE = root mean squared error.

**Table 3:** Linear regression of predictors for proportion of high effort trials selected across both task blocks.Prop. = proportion. High effort group: > 70% HR trials selected; low effort group: <= 70% HR trials selected. BDI (prop. score) = depression score on the Beck Depression Inventory, PSS =Perceived Stress Scale, NCS = Need for Cognition Scale. Block = Block of task (increasing effort block or shifting reward block). Proportion scores are scores divided by total possible score. AIC =Akaike information criterion, BIC = Bayesian information criterion, ICC = intra class correlation,RMSE = root mean squared error.

|  | Linear model: Prop. high effort trials selected |
| --- | --- |
| (Intercept) | 0.79***<br>(0.05) |
| Visual short term memory ability (K estimate) | 0.01<br>(0.01) |
| BDI (prop. score) | 0.07<br>(0.07) |
| PSS | 0.00*<br>(0.00) |
| NCS | 0.00*<br>(0.00) |
| Sex | 0.06*<br>(0.02) |
| Block | 0.00<br>(0.02) |
| Num.Obs. | 639 |
| R<sup>2</sup> | 0.042 |
| R<sup>2</sup> Adj. | 0.033 |
| RMSE | 0.22 |

+ p < 0.1, \* p < 0.05, \*\* p < 0.01, \*\*\* p < 0.001

**Table 4:** Summary table of accuracy and choice latency by group and effort level chosen. High effort group:> 70% HR trials selected; low effort group: <= 70% HR trials selected.

| Group | Effort level | Block | $M_{Accuracy}$ | $SD_{Accuracy}$ | $M_{Latency}$ | $SD_{Latency}$ |
| --- | --- | --- | --- | --- | --- | --- |
| High effort | Effort increase | high | 0.69 | 0.32 | 1.73 | 1.75 |
| High effort | Effort increase | low | 0.95 | 0.17 | 3.24 | 3.25 |
| High effort | Reward shift | high | 0.70 | 0.14 | 1.71 | 1.98 |
| High effort | Reward shift | low | 0.93 | 0.15 | 2.57 | 2.07 |
| Low effort | Effort increase | high | 0.62 | 0.36 | 1.54 | 1.44 |
| Low effort | Effort increase | low | 0.91 | 0.16 | 1.90 | 1.22 |
| Low effort | Reward shift | high | 0.68 | 0.19 | 1.43 | 1.12 |
| Low effort | Reward shift | low | 0.91 | 0.15 | 1.83 | 1.17 |

### Predictors of accuracy and choice latency

We evaluated predictors of accuracy on each trial and choice latency using k-estimate score, depression score, chronic stress score, and Need for Cognition score by sex (Fig. 5). A multi-level model analysis revealed that the effort level required in a selected trial significantly predicted accuracy across all trials such that accuracy was higher on low effort (LR) trials compared to high effort (HR) trials. Importantly, of the predictors of interest, only visual short-term memory score predicted accuracy across blocks, such that higher short-term memory ability was associated with higher accuracy (Table 5; Fig. 6). Trial effort level also predicted choice latency, with slower choices on low relative to high effort trials. K-estimate score also predicted choice latency such that higher short-term memory ability predicted slower choices across all trials (Table 5; Fig. 6). Sex, depressive levels, chronic stress scores, Need for Cognition scores, and effort preference group did not significantly predict accuracy or choice latency in either block of the task.

**Figure 5:**
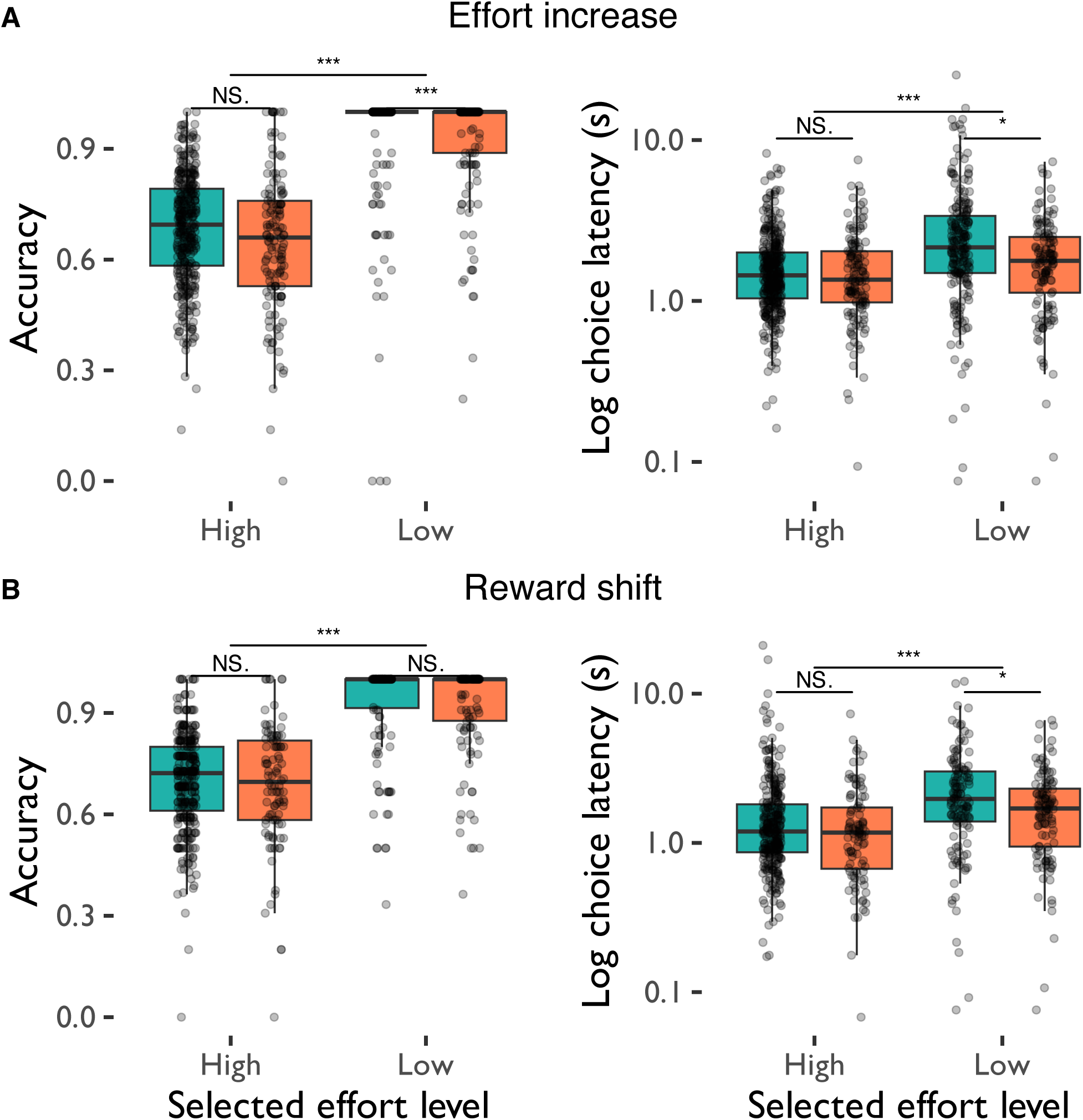
Plot of accuracy and choice latency by effort choice and effort preference group in the (A) effort increase and (B) reward shift blocks. High effort group: > 70% HR trials selected; low effort group:<= 70% HR trials selected.

**Figure 6:**
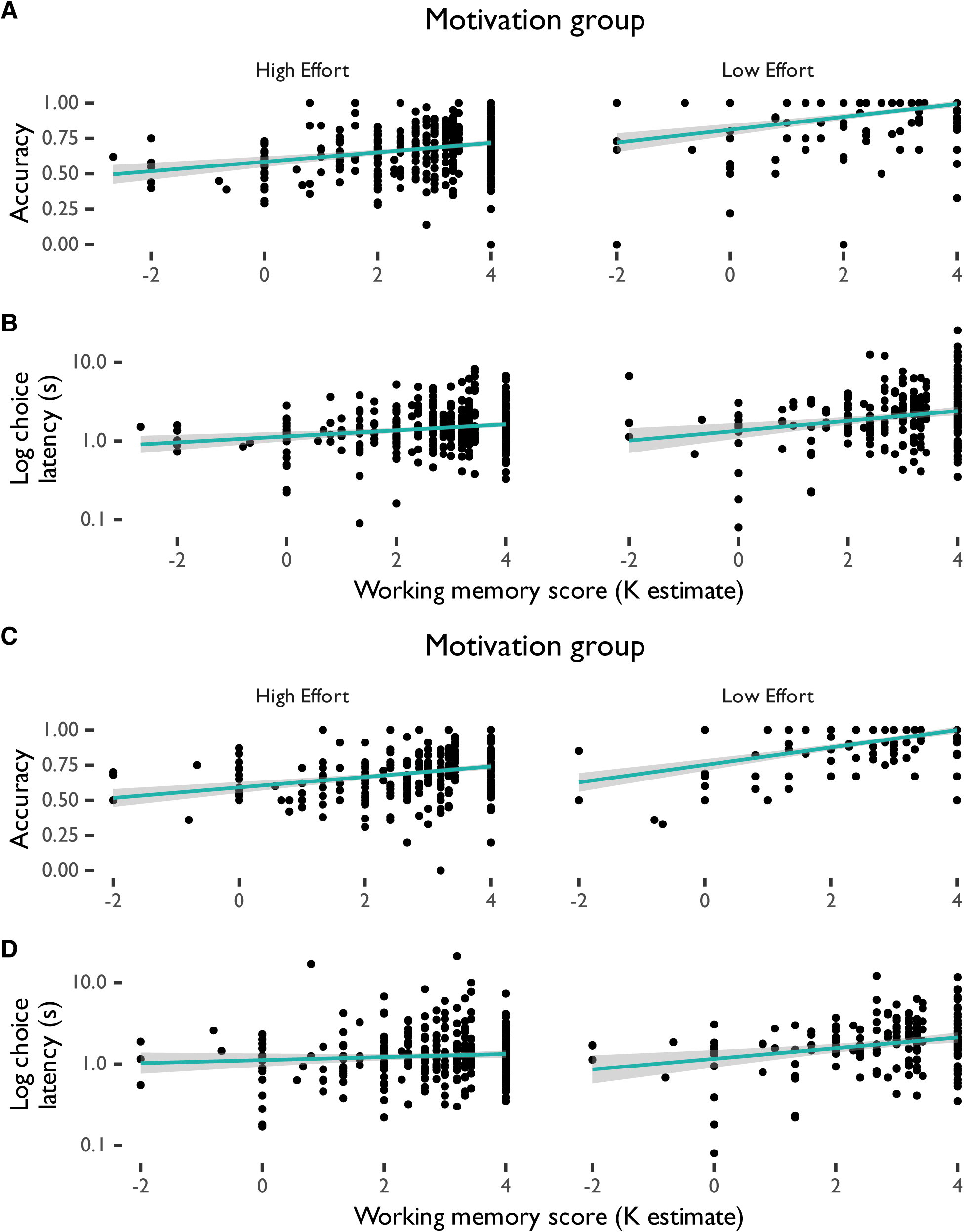
Accuracy and choice latency by visual short term memory ability. (A, C) Accuracy and (B, D) choice latency in the (A, B) shifting effort and (C, D) reward phases of the change detection task by visual short term memory ability (K estimate). The choice latency plots have been log-transformed on the y-axis to more clearly show small choice latency values.

**Table 5:**
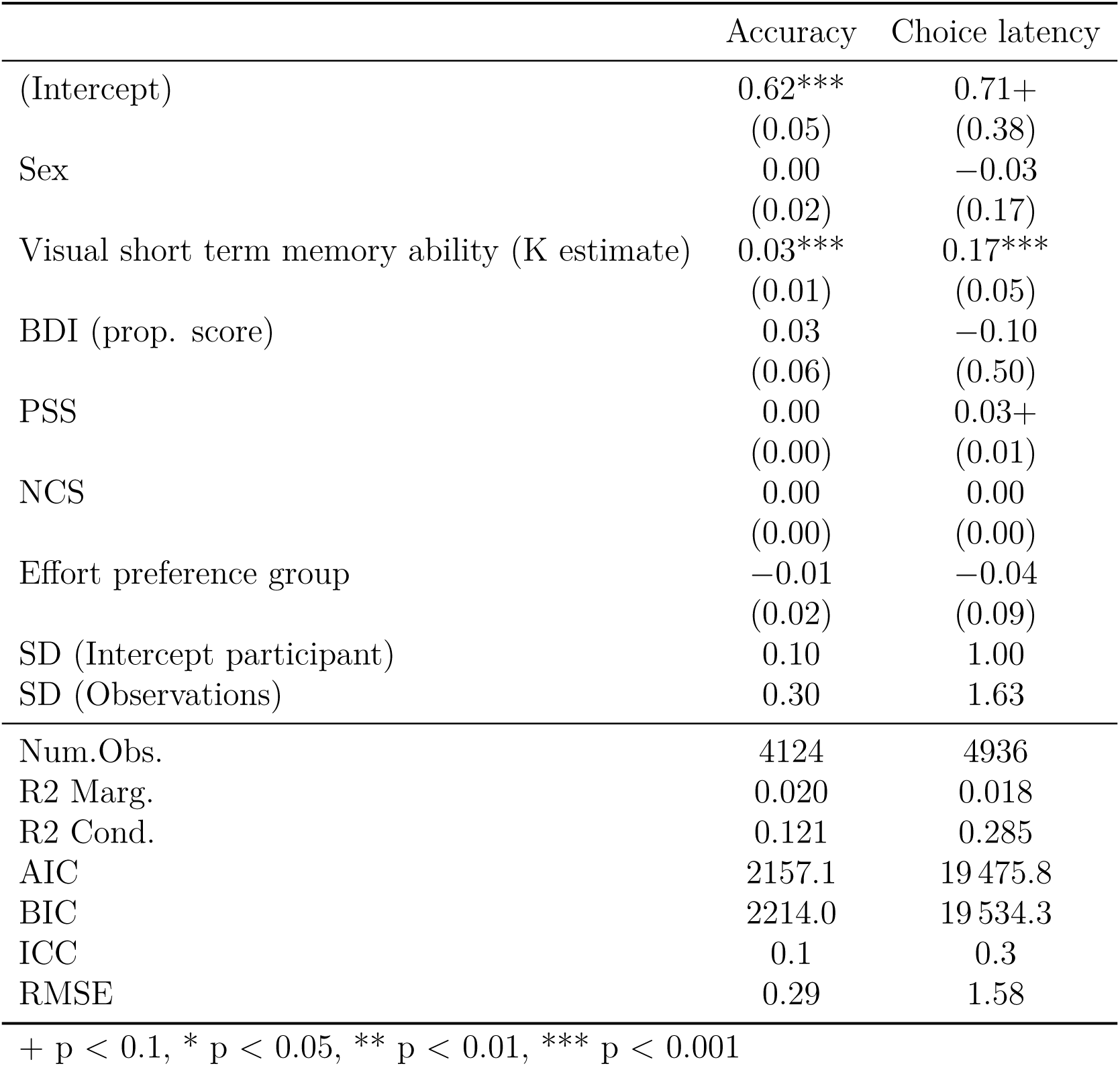
Multi-level model analysis coefficients and standard errors for accuracy and choice latency in the choice phase of the task across both effort increase and reward shift blocks. BDI (prop. score) =depression score on the Beck Depression Inventory, PSS = Perceived Stress Scale. Proportion scores are scores divided by total possible score. AIC = Akaike information criterion, BIC =Bayesian information criterion, ICC = intra class correlation, RMSE = root mean squared error.

### Equivalence in effort and reward discounting

As effort demands and reward levels were varied in each block respectively, we investigated the level of effort demands and offered reward at which participants selected high and low effort trials equally often. This point of equivalence is of interest as it indicates the point at which participants begin to discount potential rewards in favour of lower effort demands and rated both effort and reward as subjectively equivalent (Westbrook et al., 2013). We found that participants were closest to this point of equivalence in the shifting reward block when high effort trials offered a reward of 16 points (Fig. 7). However, a one-sample t-test revealed that, in the *shifting reward* block, high effort trial preferences were significantly below 50% (*t*(141) = 6.01, *p* <0.001) at this level of reward. Furthermore, high effort preferences remained significantly above 50% across all trials in the increasing effort block (*t*(554) = 23.10, *p* <0.001. Thus, participants chose low effort trials significantly more often than high effort trials when 16 points were offered in the shifting effort block. However, they selected high effort trials significantly more often than low effort trials when effort levels decreased during the task - regardless of the amount of reward offered.

**Figure 7:**
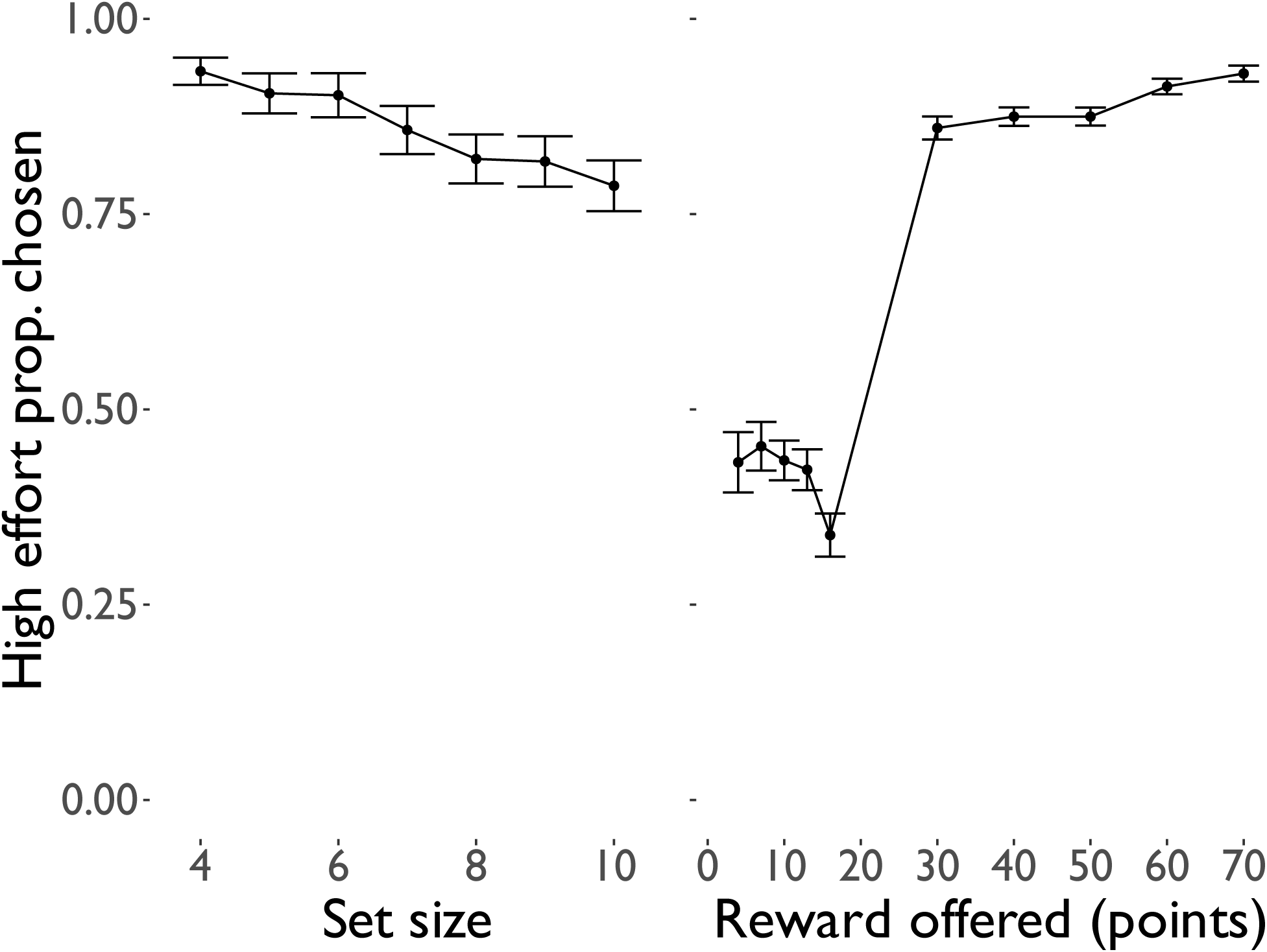
High effort preferences, expressed as a proportion of high effort trials selected out of all trials at a given effort or reward level.

### Descriptions of participant experiences

Beyond our behavioural measures of participants’ effort choices, we also asked participants to describe what effort strategies they used during the task, and whether and how their experiences of effort deployment in the task reflected real-world effort deployment decisions. We conducted an AI-assisted thematic analysis to capture open-ended groupings (Martínez-Pernía et al., 2025), as per Braun and Clarke (2006), to capture the predominant themes and sub-themes that arose from participants’ descriptions of their experiences, in order to examine what strategies participants used to complete this task vs. real-world effort tasks. We examined the proportion of participants whose responses encapsulated categories and sub-categories of themes in response to each question (Fig. 8; Table 6). From this thematic information, we found that, when discussing the effort strategies used (Q1), over 80% of participants discussed attending to the decision-making strategies they used and the intentions behind their choices, including tracking goals and maximizing rewards. When asked about their decision to select more high effort vs. low-effort trials (Q2), a majority of participants discussed their intention to seek challenges and maximize rewards, as well as effort-reward tradeoffs. When asked whether effort strategies used were comparable to those in the real world (Q3), a majority of participants discussed everyday effort strategies and their evaluations of the task’s difficulty. Furthermore, many described the in-lab task as harder, and requiring more intensive effort strategies, than real-world cognitively effortful tasks.

**Figure 8:**
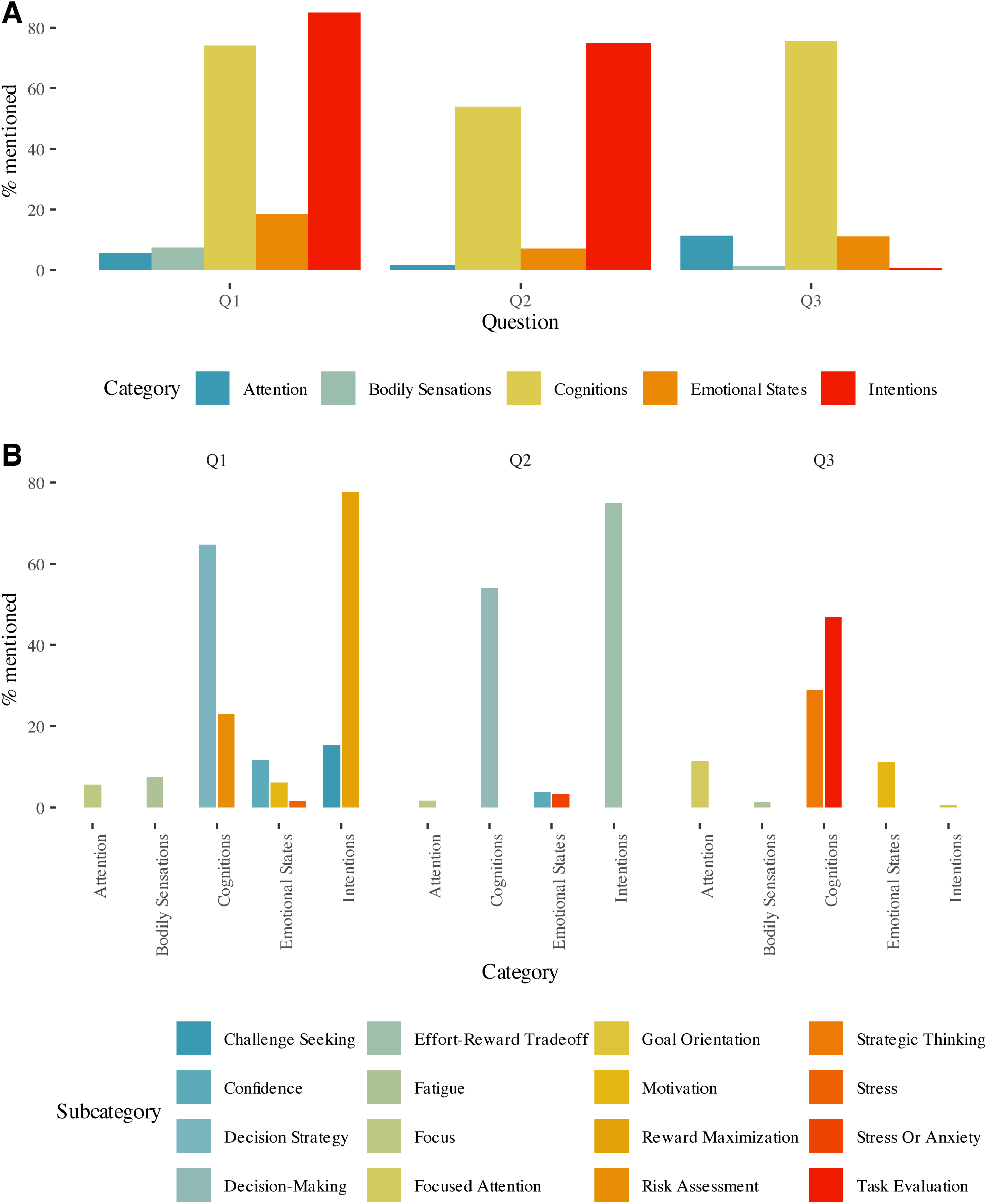
Bar plots indicating the proportion of participants whose responses captured specific A) categories and B) subcategories in responses to the following questions: Q1) “Describe what influenced your decision to choose an easy trial (less effort for a lower pay off) or a harder trial (more effort fora higher payoff) throughout the task”; Q2) “When you have to choose to use more effort to get a greater reward vs. less effort for a lesser reward in everyday life, would you make the same kind of choices you made today?”; and Q3) “To what extent do you feel that your ability to complete this task reflects your ability to complete real-world tasks, and what strategies did you use to complete this task?”

**Table 6:** Summary table of themes and sub-themes identified across participant responses following thematica nalysis.

| Category | Subcategory | Conceptual definition | Example quote |
| --- | --- | --- | --- |
| Attention | Focus | The act of concentrating on a particular task or stimulus. | "When I selected the difficult task, I would pay attention to as much as I could..." |
|  | Focused Attention | Directing mental effort toward specific stimuli or tasks. | "I think I did pretty good on this task because I used to play a lot of video games which required fast response time and high focus." |
| Bodily Sensations | Fatigue | Physical sensations of tiredness or need for rest. | "I told myself that I need a 'reset' so it's okay to do an easier one and re-orient my eyes." |
| Cognitions | Decision Strategy | The mental process of selecting a course of action among several alternatives. | "I chose the harder trials as the default, but if I felt that I was really struggling with them...I would step back and do the easy ones to...regain confidence." |
|  | Decision-Making | The participant reflects on how they make choices or evaluate options. | "It depends on the task and the reward. For example, when I'm at work and asked if I want to leave early (for less money obviously), I always choose to leave." |
|  | Risk Assessment | The evaluation of the potential risks and benefits of a decision. | "I decided to more often go for the harder trial because it is much more rewarding, and even if I mis-click, or get some wrong, there is always the next time...it felt like the best option to get more points." |
|  | Strategic Thinking | Use of deliberate methods or plans to accomplish a task. | "My strategies once I got the hang of it was to visualize the colors I saw in that general area rather than remembering the specific placement as well because that ended up distracting me...reflects how I complete real-world tasks." |
|  | Task Evaluation | Cognitive appraisal of task difficulty or performance. | "I think that I like to try doing things that are more challenging in order to not only get the large reward, but to also learn and grow from the experience." |
| Emotional States | Confidence | A feeling of self-assurance arising from one's abilities. | "I will try the easy mode first as a trial, and then, after I know I can somewhat keep up with the fast movement, I will have the confidence to try the hard one." |
|  | Motivation | Internal emotional drive to act or perform. | "When I noticed that the difficulty changed from 10 to 7 squares after a while, I would be a bit more motivated to try the harder trial again, but did not always make this decision." |
|  | Stress | A state of mental or emotional strain resulting from demanding circumstances. | "I also felt like I would be more stressed with the harder trials and I didn't want to feel that so I just did the easy trials." |
|  | Stress or Anxiety | The participant describes emotional discomfort or anxiety related to effortful tasks. | "In real life I usually get scared easily when trying new things, so I will mostly have to do a trial run/the easier mode first and then go for the harder mode." |
| Intentions | Challenge Seeking | The intention to seek out difficult tasks to test one's abilities. | "I also chose to do the harder trial sometimes as it did have a bigger reward and helped me challenge myself for personal growth." |
|  | Effort-Reward Tradeoff | The participant expresses a preference or decision-making strategy based on balancing effort and expected reward. | "If I had to put in more effort to get a greater reward, I would do it...the fact that the incentive is a lesser reward is not really a big motivation to do something." |
|  | Goal Orientation | Purposeful direction of behavior toward achieving outcomes. | "I think it shows a lot about my realistic ability to accomplish tasks, because in reality I would rather put in more effort to achieve my goals." |
|  | Reward Maximization | The intention to maximize outcomes or benefits from actions. | "When looking at the number of points compared to the easy trial and harder trial I viewed it as points = money in a 1:1 ratio." |

## Discussion

Here, we show that the personal and contextual factors that predicted participants’ choices of more lucrative, cognitively demanding options in an effort choice task depended on whether effort or reward amount were varied between trials. Specifically, we found that higher visual short term memory predicted more frequent selection of high-effort trials when reward amount - but not effort level - was manipulated. In contrast, lower perceived stress (lower PSS scores; (Levenstein et al., 1993)) and higher need for cognition (higher NCS scores; (Cacioppo and Petty, 1982)) were associated with more frequent selection of high-effort trials when effort requirements varied. Thus, stress and need for cognitive stimulation were most important when one had to evaluate effort benefits and costs, whereas cognitive ability was most important in a context eliciting sensitivity to potential rewards. The results further suggest that stress levels, cognitive capacity, and preference for cognitive challenges may be constraints that are weighed differently depending on whether reward level or effort demands are manipulated.

Critically, only visual short-term memory, as captured by higher K-estimate scores in the calibration phase of the task, predicted accuracy or choice latency during the task, similar to our previous report (Forys et al., 2024). Higher levels of perceived stress, or lower NCS scores, were thus not associated with reduced capacity to successfully completing the harder task. If more stressed individuals were less able to complete harder trials - or those who enjoy cognitively demanding tasks were better at performing them - it would be rational to select the easier option. Instead, these factors can capture choices to exert cognitive effort for more reward, independent of one’s ability to successfully attain those rewards.

Furthermore, this non-normative behaviour is only seen when effort costs are varied, suggesting differential processing of shifting vs. static effort costs. This conclusion is further supported by our finding that working memory capacity did not predict choice of high effort trials when effort costs shift.

Our results contextualize work by Westbrook et al. (2013), Crawford et al. (2021), and Westbrook et al. (2019) showing that low preferences for cognitively effortful strategies are associated with greater effort discounting and increased low effort trial preferences over time. In our study, we varied effort demands and reward levels across both effort options, while Westbrook et al. (2013) focused on varying reward levels across stable effort choices. Westbrook et al. (2013) show that increased high effort preferences are associated with increased NCS scores. Thus, varying rewards based on choices may invite increased engagement with everyday effort strategies. The general pattern of effort preferences observed in our study also aligns with that found in rodents in the rCET (Hosking et al., 2016; Hales et al., 2024). As with humans in our task, rodents that chose more higher effort trials selected them faster. Thus, our study elucidates translationally relevant effects specific to tracking changing reward preferences and effort demands.

Past findings from Kramer et al. (2021) show increased willingness to deploy cognitive effort given higher NCS scores and cognitive capacity (using Raven’s progressive matrices; Hamel and Schmittmann (2006)). Depending on the nature of the task demands presented (e.g., whether the task engages visuospatial or linear memory demands), awareness of one’s cognitive capacity could overlap with preferences for choosing more cognitively effortful strategies (Lopez-Gamundi and Wardle, 2018). A caveat is that choices to deploy more effort for the same level of reward may not be captured by general effort deployment ability if fatigue dampens one’s motivation to complete the task (Inzlicht et al., 2018). However, work comparing effects of fatigue on paired cognitive vs. physical effort paradigms shows significant effects of fatigue on task performance for physical but not cognitive effort (Lopez-Gamundi and Wardle, 2018).

Another potential contributor to the observed results is whether participants can correctly forecast their inability to achieve their desired outcomes in the task. For example, if participants are not confident in their abilities in the task (Chung et al., 2024), they may not be able to learn effectively from negative outcomes, continue to use suboptimal effort deployment strategies, and fail to adapt to changing levels of fatigue. To this point, McIntosh et al. (2021) found that the ability to forecast both positive and negative outcomes when generating ideas improved the likelihood that these forecasts would be correct. Increased levels of chronic stress could thus be associated with fewer choices of high effort trials through impairments to effective effort forecasting, along with more rapid delay discounting (Massar et al., 2020).

Laboratory-based decision making tasks present simplified models of real-world choices (Bustamante et al., 2023; Trier et al., 2025), and we must ask whether our task accurately captures naturalistic behaviours. To this end, we also asked participants to describe whether task demands reflected real-world cognitive effort decisions. They reported choosing more high-effort trials if they felt that they could effectively memorize the configuration of the shapes, and attended to task difficulty when considering what strategies to use to complete the task. Additionally, many participants reported that they were willing to challenge themselves to select high-effort trials given higher payoffs. Some participants felt that the task reflected real-world cognitive effort demands, but was more challenging than what they would face outside the lab. Through an AI-assisted thematic analysis

(Martínez-Pernía et al., 2025), we further identified several key themes and sub-themes that characterized participant experiences in the task. These included the finding that participants focused on maximizing rewards while managing stress level; tracking their confidence in their ability to make decisions; and considering whether and when to trade off further rewards given excessive effort demands. The themes that participants most frequently discuss as being relevant to in-task and everyday effort choices converge with factors that predicted patterns of cognitive effort decision-making, including the association between cognitive effort capacity and willingness to choose more high-effort trials. This thematic analysis complements the behavioural findings of our task to indicate their relevance to out-of-lab scenarios where participants are in constant engagement with - and impacting - a dynamic environment, not merely responding to stimuli. Furthermore, our work represents one of the first applications of large language models to large-scale textual analysis tasks in a cognitive science context, holding promise for future use of this technology to aid researchers in rapidly making sense of extensive subjective report datasets.

Importantly, willingness to deploy more effort for the same level of reward may not be captured by an overall measure of effort deployment ability if fatigue dampens motivation to complete it (Inzlicht et al., 2018). However, past work shows significant effects of fatigue on task performance for physical but not cognitive effort (Lopez-Gamundi and Wardle, 2018). Furthermore, increased chronic stress, as captured by the PSS (Levenstein et al., 1993), could be associated with excessive exploitation of present, sub-optimal effort strategies rather than seeking out new, more optimal ones (Lenow et al., 2017; Bogdanov et al., 2021). Findings on our visual short-term memory paradigm further suggest that increasing effort demands could also increase suboptimal decision-making strategies given high levels of chronic stress.

We must note limitations qualifying our results. First, there was no convergence in the level of high vs. low effort trial rewards in the shifting reward block. Thus, we could not directly compare choices for high vs. low effort trials at the same level of reward (Westbrook et al., 2013). However, a point of equivalence was approached and, thus, an overall evaluation of subjective equivalence was still possible (Fig. 7). Future studies should include reward convergence between high and low effort trials, to determine if this is the point at which subjective equivalence (equal preference for low and high effort trials) is achieved. Second, effort choices could be explained by willingness to deploy effort given decreasing rewards vs. ability to meet increasing effort demands. For example, the same effort level may feel more fatiguing if the reward on offer decreases - and people may disengage from deploying more cognitive effort as effort demands increase because they are unwilling to deploy more effort to get the same reward. Thus, in future studies, one should directly manipulate the correspondence between effort required and rewards available on high vs. low effort trials to disambiguate factors impacting one’s willingness vs. ability to deploy effort. Third, outcomes in the task were exclusively framed as rewards and not as losses. Loss avoidance may motivate cognitive effort deployment more strongly reward-seeking (Massar et al., 2016). Thus, future studies should also evaluate the impact of loss aversion on one’s motivation to select high vs. low effort trials. Last, we did not directly manipulate preferences for cognitive capacity, effort strategies, or stress levels in our task, limiting the causal inferences we can draw from our results. Future work should incorporate framing tasks to prime participants’ expectations of the amount of information they should attend to in a cognitively effortful tasks.

Overall, our findings suggest a differential weighing of personal factors constraining the choice to expend effort for reward. Levels of cognitive capacity, indexed as short-term working memory ability, predicted preferences for more cognitively effortful options given increasing effort demands. Individual differences in experience, such as chronic stress, and traits such as enjoyment of cognitive effort, predicted participants’ choices to engage with increasingly difficult cognitive effort demands to obtain greater benefits. Self-report descriptions of participants’ experiences suggested that our task’s demands, and strategies used to make choices in the task, were generally relevant to everyday situations. These results extend past work, including rodent studies, to shed light on nuances of the traits and experiences that capture everyday choices about cognitive effort. Furthermore, they suggest areas of interest when considering clinical interventions to target chronic stress through the reweighing of attention towards rewards away from effort costs. They contribute to an emerging picture of real-world effortful decision making as highly dynamic and context-dependent, modulated by variable weightings of multiple predictive factors.

## Data and code availability

All task code, stimuli, data and code used to generate this manuscript and the figures are available at https://osf.io/x2nwc/.

